# Stressed mothers, tolerant daughters: a case study about the physiological responses and growth of sugarcane plants under water deficit

**DOI:** 10.1101/448241

**Authors:** Fernanda C. C. Marcos, Neidiquele M. Silveira, Paulo E. R. Marchiori, Eduardo C. Machado, Gustavo M. Souza, Marcos G. A. Landell, Rafael V. Ribeiro

**Affiliations:** Laboratory of Crop Physiology, Department of Plant Biology, Institute of Biology, University of Campinas (UNICAMP), Campinas SP, Brazil; Laboratory of Plant Physiology ‘Coaracy M. Franco’, Centre for Research and Development in Ecophysiology and Biophysics, Agronomic Institute (IAC), Campinas SP, Brazil; Department of Biology, Federal University of Lavras (UFLA), Lavras MG, Brazil.; Department of Botany, Institute of Biology, Federal University of Pelotas (UFPel), Pelotas RS, Brazil.; Sugarcane Research Center, IAC, Ribeirão Preto SP, Brazil.

**Keywords:** drought, memory, photosynthesis, propagation, resistance.

## Abstract

Drought stress can imprint marks in plants after a previous exposure, leading to a permissive state that facilitates a more effective response to subsequent stress events. Such stress imprints would benefit plants obtained from progenitors previously exposed to drought. Herein, our hypothesis was that daughter plants obtained from mother plants previously exposed to water deficit will perform better under water deficit as compared to those obtained from mothers that did not face stressful conditions. Sugarcane mother plants were grown under well-hydrated conditions or subjected to three cycles of water deficit by water withholding. Then, daughter plants produced through vegetative propagation were subjected to water deficit. Leaf gas exchange was reduced under water deficit and daughters from mothers that experienced water deficit presented a faster recovery of CO_2_ assimilation and higher instantaneous carboxylation efficiency after rehydration as compared to daughters from mothers that did not face water deficit. Plants obtained from mother plants that faced water deficit showed the highest leaf proline concentration under water deficit as well as higher leaf H_2_O _2_ concentration and leaf ascorbate peroxidase activity regardless of water regime. Under well-watered conditions, daughters from mothers that faced stressful conditions presented higher root H2O2 concentration and root catalase activity than ones from mothers that did not experience water shortage. Such physiological changes were associated with improvements in leaf area and shoot and root dry matter accumulation in daughters from stressed mothers. Our results suggest that root H_2_O_2_ concentration is a chemical signal associated with stress memory and improved sugarcane growth. Such findings bring a new perspective to sugarcane production systems, in which stress memory can be explored for improving drought tolerance in rainfed areas.

## Introduction

As a semi-perennial species, sugarcane plants face seasonal drought under field conditions, where water deficit causes reduction in photosynthesis and accumulation of carbohydrates, changes in antioxidant metabolism, and finally impairment of plant growth and sucrose yield [1, 2]. However, recurrent cycles of drought followed by rehydration are known to improve plant performance during a new stressful event [3-5]. Such phenomenon indicates that plants are able to change their metabolism and growth after an external stimulus, improving recovery of photosynthesis, increasing intrinsic water use efficiency [4] and photoprotection [6] and reducing the negative impact of drought on yield [7].

Improved plant response induced by previous exposure to a limiting factor is an evidence of stress memory, a way to storage information of stressful events [3, 8, 9]. In fact, such stress memory can assist plants in future stresses [9] and one important issue is the site (plant tissue) in which information is stored within plants. Plants do not have a specific region to store information and they can sense the environment with all their body and the intricate cell signaling system. Then, plants can perceive one stimulus in one site and the respective response be found in a different organ due to signaling [10]. One important requirement for retaining information is that stress-induced signals are still present when the stressor is no longer affecting plants [11].

In nature, plant phenotype is also defined by transgenerational regulation, which occurs when internal changes persist in the next generation through epigenetic marks such as DNA methylation [8]. There is reasonable evidence for assuming that plants can sense changes in the environment during growth and modify the phenotype of their progeny to be more adapted to growing conditions [8, 12]. The stress-induced memory can be transferred to subsequent generations by seeds and through vegetative propagation [13]. In the first case, plants can pass epigenetic information through the meiosis process and produce seeds with stress memory [12, 14]. For instance, Boyko et al. [14] showed that *Arabidopsis thaliana* exposed to cold, heat and flooding had increased global genome methylation and higher tolerance to stress as compared to progeny from plants that never faced stressful conditions. However, stress-induced signals may be erased or diminished during meiosis, reducing stress memory. On the other hand, clonal plants produced by vegetative propagation have apparently better ability to recover signals acquired during stress events than non-clonal plants [15].

Considering stress memory, plant propagation and drought-induced effects on plants, we hypothesized that plants obtained from others previously exposed to drought will perform better under water deficit as compared to plants obtained from mother plants that never faced water shortage. Through vegetative propagation, information about previous stresses (memory) can be stored in sugarcane buds, which will sprout and produce new plants. Sugarcane is an important crop for ethanol and bioenergy production – a clean alternative for energy production – and its expansion to rainfed areas needs more drought tolerant plants. Then, stress memory would be an interesting tool for improving crop establishment and initial growth in such new areas.

## Materials and methods

### Plant material and growth conditions

Sugarcane *(Saccharum* spp.) plants cv. IACSP94-2094 were obtained from mini-stalks containing one bud and grown in plastic pots (0.5 L), with commercial substrate composed of sphagnum peat, expanded vermiculite, limestone dolomite, agricultural gypsum and NPK fertilizer (Carolina Soil®, Vera Cruz RS, Brazil). Thirty-four days after planting, plants were transferred to larger pots (20 L) containing typical red-yellow Latosoil [16], fertilized with urea (equivalent to 300 kg N ha^−1^), superphosphate (equivalent to 300 kg P_2_O_5_ ha^−1^) and potassium chloride (equivalent to 260 kg K_2_O ha^−1^) according to Dias and Rossetto [17]. During the experiment, other three fertilizations were performed at 30, 60 and 150 days after planting, with the same amount of urea, superphosphate and potassium chloride as the first fertilization. The plants were grown under greenhouse conditions, where the average air temperature was 24.4±6.6 °C, relative humidity was 76±17% and the maximum photosynthetic photon flux density (PPFD) was approximately 1,200 µmol m^−2^ s^−1^. Plants were irrigated daily and grown under well-hydrated conditions until they were six-month old.

### Inducing water deficit to mother plants

When plants were 6-month old, one group of plants was maintained under daily irrigation (W) and another group was subjected to three cycles of water deficit (D) by water withholding. Each cycle of water deficit lasted nine days and soil moisture was monitored with soil moisture-sensors model Water Scout SM100 (Yara ZimTechnology, Berlin, Germany). While soil volumetric water content (VWC) reached 20% during cycles of water deficit, it was higher than 60% in well-watered pots. After nine days of water deficit, plants were irrigated and maintained under well-watered conditions for six days before the new cycle of water deficit, with leaf gas exchange being measured daily. After three cycles of water deficit, we evaluated the number of tillers, number of green and senescent leaves, total leaf area and dry matter of leaves, stems and roots. Then, daughter plants were produced through vegetative propagation from those mother plants that experienced or not cycles of water deficit, as cited in the previous section.

### Inducing water deficit to daughter plants

After sprouting in commercial substrate (Carolina Soil®, Vera Cruz RS, Brazil), one-month old plants were placed in plastic boxes (12 L) with nutrient solution and transferred to a growth chamber (PGR15, Conviron, Winnipeg MB, Canada) under air temperature of 30/20 °C (day/night), with 12 h photoperiod, air relative humidity of 80% and PPFD of 800 µmol m^−2^ s^−1^. Only the root system was immersed in modified Sarruge [18] nutrient solution (15 mmol L^−1^ N [7% as NH_4_+]; 4.8 mmol L^−1^ K; 5.0 mmol L^−1^ Ca; 2.0 mmol L^−1^ Mg; 1.0 mmol L^−1^ P; 1.2 mmol L^−1^ S; 28.0 µmol L^−1^ B; 54.0 µmol L^−1^ Fe; 5.5 µmol L^−1^ Mn; 2.1 µmol L^−1^ Zn; 1.1 µmol L^−1^ Cu and 0.01 µmol L^−1^ Mo). Nutrient solution was renewed in week intervals and pH was maintained at 5.8±0.2 and electrical conductivity at 1.72±0.18 mS cm^−1^. The osmotic potential of nutrient solution was −0.12 MPa. Two boxes containing plants obtained from irrigated mother plants and two boxes containing plants from those mothers subjected to three cycles of water deficit were prepared.

Forty-eight days after transferring plants to the hydroponic system, one group of plants was subjected to water deficit by adding PEG-8000 (CarbowaxTM PEG-8000, Dow Chemical Comp, Midland MI, USA) to the nutrient solution for nine days. We added PEG-8000 gradually to prevent osmotic shock. Then, the osmotic potential of nutrient solution was reduced to −0.27, −0.57 and −0.77 MPa in three consecutive days. After nine days, the plants were recovered by supplying them with a nutrient solution with osmotic potential of −0.12 MPa (control condition) for five days. At the end, four treatments were defined taking into account the plant origin and also the water regime plants were facing: plants obtained from mother plants grown under well-watered conditions and then maintained under well-watered conditions (W/W); plants obtained from mother plants grown under well-watered conditions and then subjected to water (W/D); plants obtained from mother plants that experienced water deficit and then maintained under well-watered conditions (D/W); plants obtained from mother plants that faced water deficit and then subjected to water (D/D).

### Leaf gas exchange and photochemistry

Leaf gas exchange was measured daily with an infrared gas analyzer (LI-6400, LICOR, Lincoln NE, USA) attached to a modulated fluorometer (6400-40 LCF, LICOR, Lincoln NE, USA). The measurements were performed between 10:00 and 13:00 h under PPFD of 2,000 µmol m^−2^ s^−1^ and air CO_2_ concentration of 380 µmol mol^−1^. CO_2_ assimilation *(A),* stomatal conductance (g_S_), intercellular CO_2_ concentration (C_i_), transpiration (E), intrinsic water use efficiency (A/g_S_), and the instantaneous carboxylation efficiency (*k=A/C*_i_) were evaluated in fully expanded leaves. *A* and *E* values were integrated throughout the experimental period to estimate the total CO_2_ gain (A_i_), the total H_2_O loss through transpiration (E_i_, and the integrated water use efficiency (WUE=A_i_/E_i_). The integrated values were estimated assuming that the values measured between 10:00 and 13:00 h were constant during the 12 hours of photoperiod. Chlorophyll fluorescence was measured simultaneously to leaf gas exchange and the apparent electron transport rate (ETR) was estimated as ETR=Φ_PSII_× PPFD × 0.85 × 0.4, in which Φ_PSII_ is the effective quantum efficiency of photosystem II (PSII), 0.85 is the light absorption and 0.4 is the fraction of light energy partitioned to PSII [19, 20]. Additionally, the non-photochemical quenching of fluorescence (NPQ) was evaluated and ETR/*A* calculated. In leaf tissues adapted to darkness (30 min), the potential quantum efficiency of photosystem II (*F*_V_/*F*_M_) was estimated [20].

### Leaf water potential and relative water content

Leaf water potential (Ψ) was evaluated at the predawn with a pressure chamber (model 3005, Soilmoisture Equipment Corp., Santa Barbara CA, USA). The leaf relative water content RWC) was calculated using the fresh (FW), turgid (TW) and dry (DW) weights of leaf discs according to Weatherley [21]: RWC = 100 × (FW − DW)/(TW − DW). Both variables were measured in daughter plants at the maximum stress condition (9^th^ day of water deficit) and recovery period.

### Carbohydrates and proline

The extraction of total soluble carbohydrates (SS) was done with methanol:chloroform:water solution [22] and quantified by the phenol–sulfuric acid method [23]. Sucrose content was quantified following van Handel [24] and starch (Sta) was determined by the enzymatic method proposed by Amaral et al. [25]. The concentration of nonstructural carbohydrates (NSC) in leaves and roots was calculated as NSC=SS+Sta. Total NSC was calculated considering the dry matter of each plant (mg plant^−1^). Plant nonstructural carbohydrates were calculated by the sum of leaf and root carbohydrates and carbohydrate partitioning among sugar types was also evaluated in both organs.

Leaf proline content was determined in test tubes by the reaction with the ninhydrin reagent (ninhydrin, acetic acid and orthophosphoric acid), glycine and acetic acid for 35 minutes at 100°C. The reaction mixture was extracted with toluene and the proline concentration was determined from a standard curve [26].

### Hydrogen peroxide and antioxidant enzymes

Evaluation of hydrogen peroxide (H_2_O_2_) was performed in 0.16 g fresh tissue (leaves and roots) ground in liquid nitrogen with the addition of polyvinylpolypyrrolidone (PVPP) and 0.1% of trichloroacetic acid (TCA) solution (w/v) [27]. The extract was centrifuged at 12,000 *g,* 4°C for 15 min. The crude extract was added to the reaction medium (1.2 mL of KI 1 mol L^−1^, potassium phosphate buffer pH 7.5 and 0.1 mol L^−1^) in microtubes and incubated on ice under dark for 1 h. After this period, the absorbance was evaluated at 390 nm. The calibration curve was done with H2O2 and results were expressed as µmol g^−1^ FW.

Enzymes were extracted from 0.2 g of fresh tissues of leaves and roots grounded in liquid nitrogen, with 1% of PVPP and 2 mL of extraction medium composed by 0.1 mol L^−1^ potassium phosphate buffer (pH 6.8), 0.1 mmol L^−1^ ethylenediaminetetraacetic (EDTA) and 1 mmol L^−1^ phenylmethylsulfonyl fluoride (PMSF). This homogenate was centrifuged at 15,000 *g* for 15 min and 4°C and the supernatant was collected and preserved on ice.

Superoxide dismutase (SOD, EC 1.15.1.1) activity was evaluated in a reaction medium with 3 mL of 100 mmol L^−1^ sodium phosphate buffer (pH 7.8), 50 mmol L^−1^methionine, 5 mmol L^−1^ EDTA, deionized water, crude extract, 100 µmol L^−1^ riboflavin and 1 mmol L^−1^ nitro blue tetrazolium chloride (NBT). A group of tubes was exposed to light (fluorescent lamp of 30 W) for 15 min, and another group remained in darkness. The absorbance was measured at 560 nm and one unit of SOD is the amount of enzyme required to inhibit the NBT photoreduction in 50% [28]. SOD was expressed as U g^−1^ FW min^−1^.

Catalase (CAT, EC 1.11.1.6) activity was assayed in a reaction medium of 3 mL of 100 mmol L^−1^ potassium phosphate buffer (pH 6.8), deionized water, 125 mmol L^−1^ H_2_O _2_ and crude extract. The decrease in absorbance at 240 nm was measured and CAT activity was estimated using a molar extinction coefficient of 36 M^−1^ cm^−1^ and expressed as nmol g^−1^ FW min^−1^ [29].

For ascorbate peroxidase (APX, EC 1.11.1.11) activity, the reaction medium was composed by 3 mL of 100 mmol L^−1^ potassium phosphate buffer (pH 6.0), deionized water, 10 mmol L^−1^ ascorbic acid, 10 mmol L^−1^ H_2_O_2_ and crude extract. The decrease in absorbance at 290 nm was measure and we used a molar extinction coefficient of 2.8 M^−1^ cm^−1^ to estimate APX in nmol g^−1^ FW min^−1^ [30].

### Biometry

The total leaf area was measured using the LI-3000 leaf area meter (LICOR, Lincoln NE, USA), and shoot and root dry matter were evaluated after drying samples in a forced air oven at 65 °C. Measurements were taken at the end of the experimental period.

### Statistical analysis

The experimental design was in randomized blocks and the causes of variation were water conditions (two levels) and material origin (two levels). The data were subjected to ANOVA procedure and the mean values (n=4) were compared by the Tukey test at 5% probability level.

## Results

### Mother plants under water deficit

Herein, mother plants are defined as those ones that provided vegetative material for propagation, i.e., small stalk segments with buds. Mother plants were subjected to three cycles of water deficit and leaf gas exchange was measured during dehydration and rehydration stages (S1 Fig). There was a significant reduction in leaf CO_2_ assimilation after four days of water withholding in all cycles of water deficit (S1 Fig), with net photosynthesis reaching null values or even negative ones (respiration). Full recovery of leaf CO_2_ assimilation was noticed in all cycles and the negative impact of water deficit was reduced from the first to the third cycle (S1 Fig). After three cycles of water deficit, there was a significant reduction of biomass production (S2 Fig), with reductions in the number, dry matter and area of green leaves as well as decreases in root and stem dry matter (S1 Table).

Then, small stalk segments (around 3 cm) with one bud were obtained from those mother plants and planted in individual recipients to produce new plants, i.e., daughter plants. Buds from mother plants subjected to water deficit had higher sprouting (ca. 95%) than buds from mother plants maintained under well-watered conditions *(ca.* 74%). Thirty days after planting, daughter plants were placed in plastic boxes with nutrient solution and four treatments were done after 18 days: plants from mother plants grown under well-watered conditions maintained under well-watered conditions (W/W) or subjected to water deficit (W/D); and plants from mother plants grown under cycles of water deficit maintained under well-watered conditions (D/W) or subjected to water deficit (D/D).

### Daughter plants under water deficit

Water deficit reduced leaf CO_2_ assimilation, stomatal conductance and the instantaneous carboxylation efficiency, regardless of the plant origin (Fig 1). Interestingly, plants originated from mother plants that experienced water deficit (D/D) presented a faster recovery of leaf CO_2_ assimilation and carboxylation efficiency as compared to W/D plants (Fig 1A,C). Integrated leaf CO_2_ assimilation and transpiration were reduced by water deficit in a similar way when comparing W/D and D/D treatments (Fig 2A,B). However, recovery of photosynthesis was favored in D/D plants and then integrated water use efficiency was improved under water deficit (Fig 2C).

**Fig 1.**
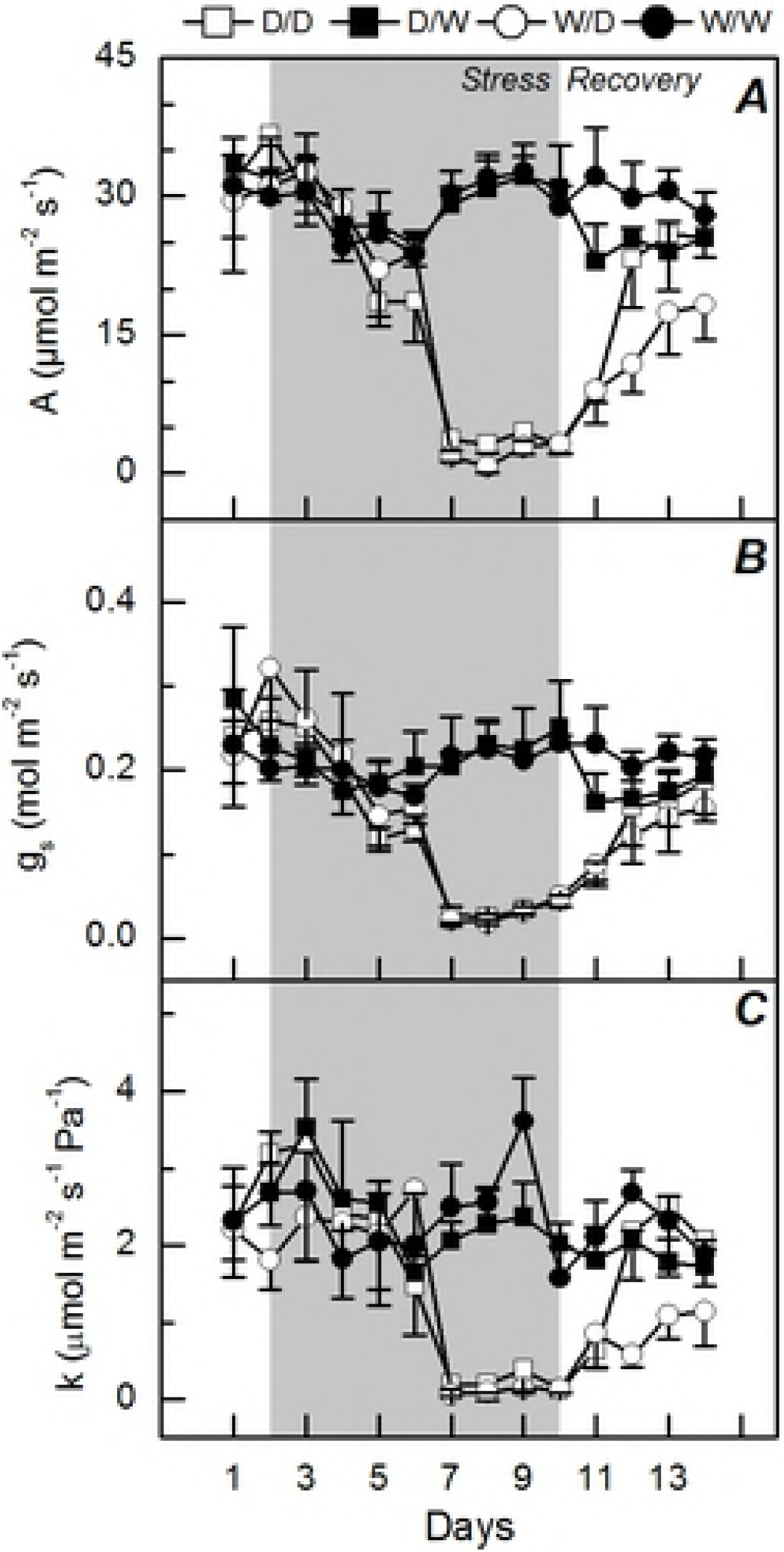
Time course of leaf gas exchange in daughter plants under water deficit. Leaf CO_2_ assimilation (*A*), stomatal conductance (*B*), and instantaneous carboxylation efficiency (*C*) in sugarcane plants grown under well-watered conditions (W/W and D/W) or subjected to water deficit (W/D and D/D). Daughter plants were obtained from mother plants previously exposed to water deficit (D/W and D/D) or grown under well-watered conditions (W/W and W/D). The gray area indicates the water deficit period. Each symbol is the mean values ± s.d. (n=4).

**Fig 2.**
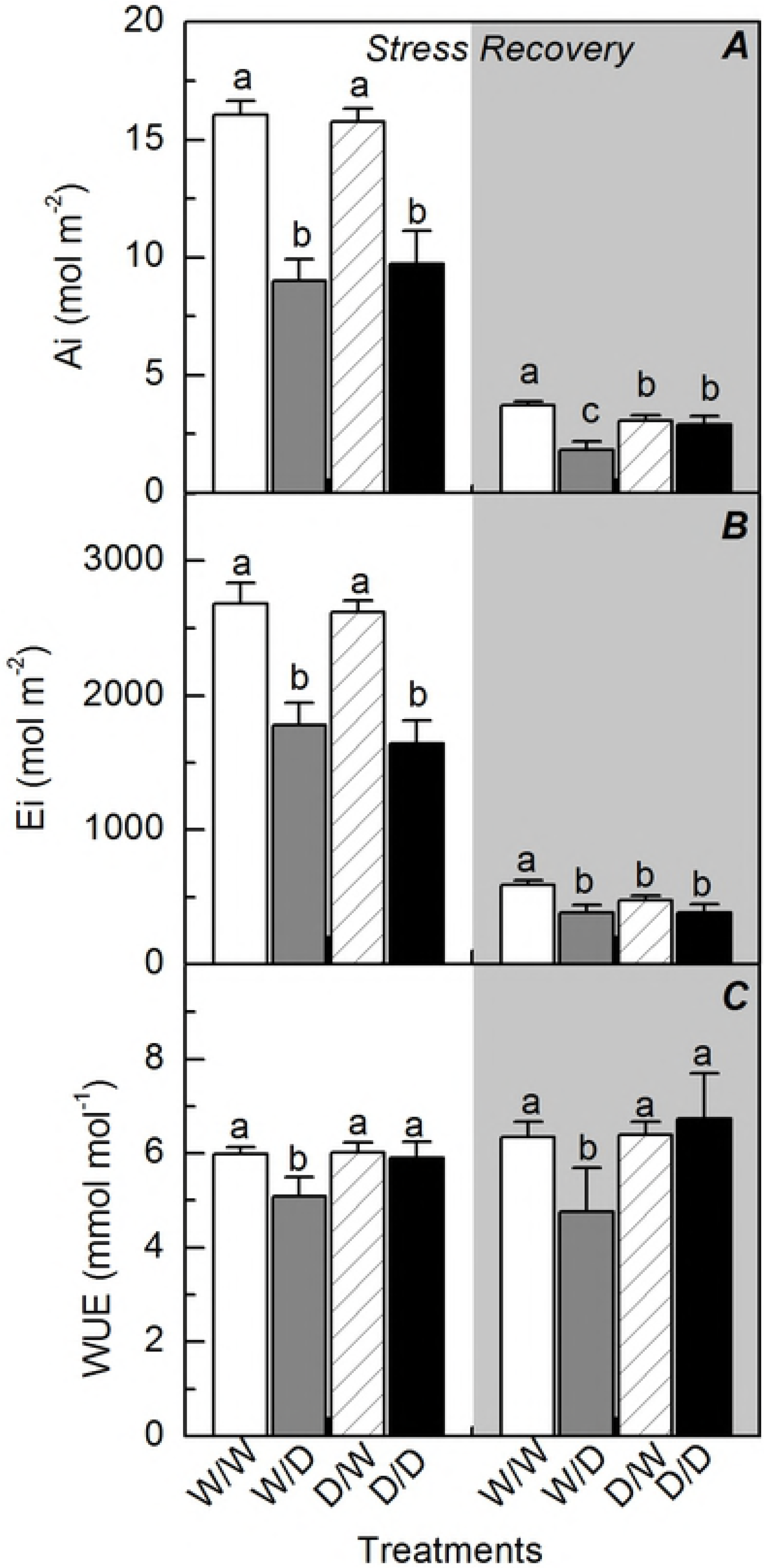
Integrated leaf gas exchange in daughter plants during and after water deficit. Integrated CO_2_ assimilation (*A*), transpiration (*B*) and water use efficiency (*C*) in sugarcane plants grown under well-watered conditions (W/W and D/W) or subjected to water deficit (W/D and D/D). Daughter plants were obtained from mother plants previously exposed to water deficit (D/W and D/D) or grown under well-watered conditions (W/W and W/D). Integration was done during the water deficit (stress) and recovery (gray area) periods, as shown in Fig 1. Each histogram is the mean values + s.d. (n=4). Different letters mean statistical differences among treatments (p<0.05).

After nine days of water deficit, pre-dawn leaf water potential was reduced and D/D plants showed the lowest values (Fig 3A). Regarding the leaf relative water content, there was a similar response to water deficit and both W/D and D/D plants exhibited the lowest values (Fig 3B). While the pre-dawn leaf water potential was fully recovered, leaf relative water content was partially recovered after four days of plant rehydration (Fig 3).

**Fig 3.**
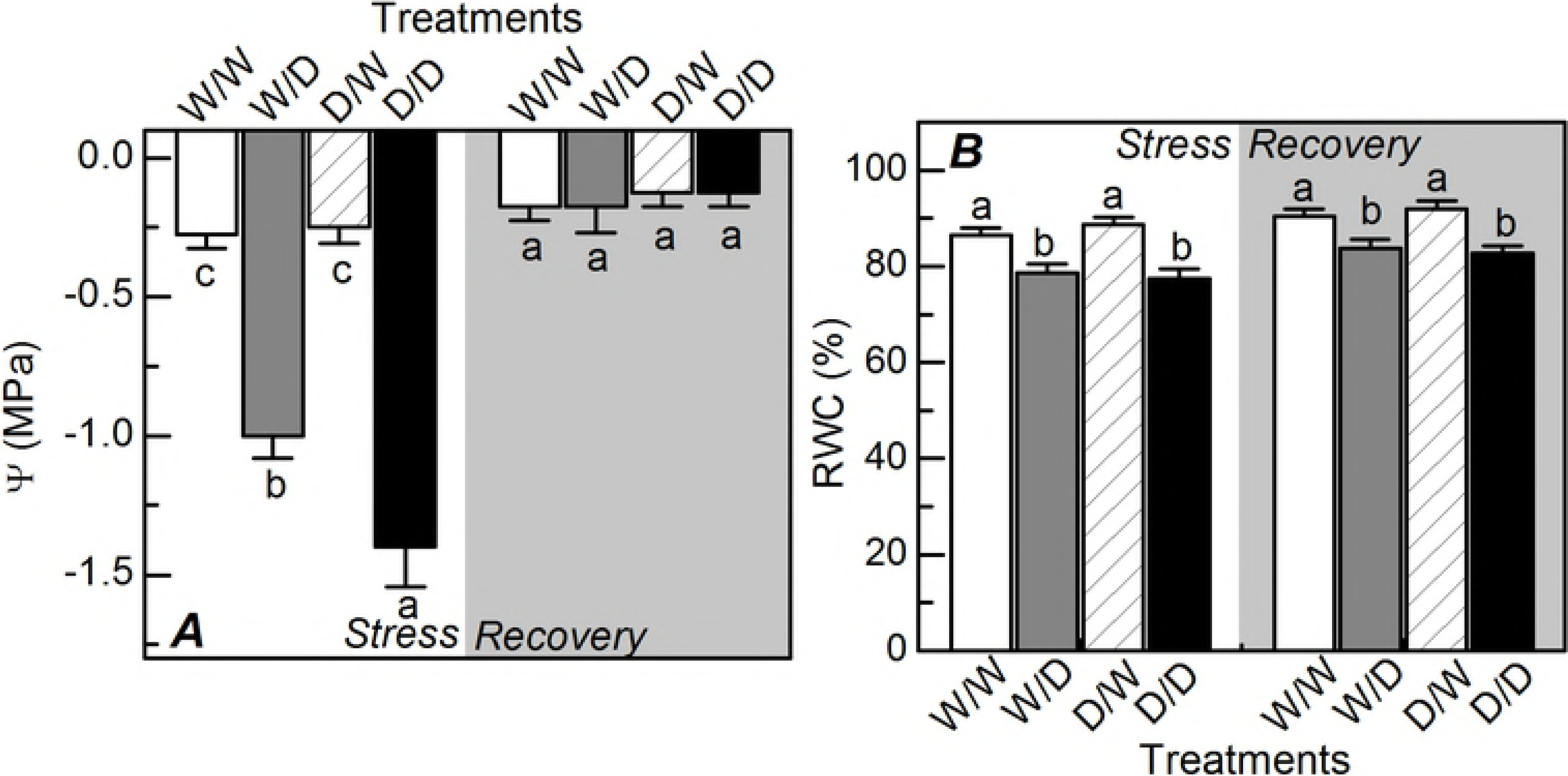
Water relations in daughter plants under water deficit. Predawn leaf water potential (*A*) and relative water content (*B*) in sugarcane plants grown under well-watered conditions (W/W and D/W) or subjected to water deficit (W/D and D/D). Daughter plants were obtained from mother plants previously exposed to water deficit (D/W and D/D) or grown under well-watered conditions (W/W and W/D). Measurements were done during the water deficit (stress) and recovery (gray area) periods, as shown in Fig 1. Each histogram is the mean values + s.d. (n=4). Different letters mean statistical differences among treatments (p<0.05).

Water deficit caused decreases in the potential quantum efficiency of PSII (F_v_/F_m_) and also in the apparent electron transport rate (ETR) of W/D and D/D plants (Fig 4A,B). Although D/D plants had shown the lowest ETR values, the ratio ETR/A was similar between W/D and D/D plants, increasing in more than three times due to water deficit (Fig 4C). Non-photochemical quenching was increased by water deficit only in W/D plants (Fig 4D). All photochemical indices were recovered after plant rehydration, with W/W vs. W/D and D/W vs. D/D plants showing similar values.

**Fig 4.**
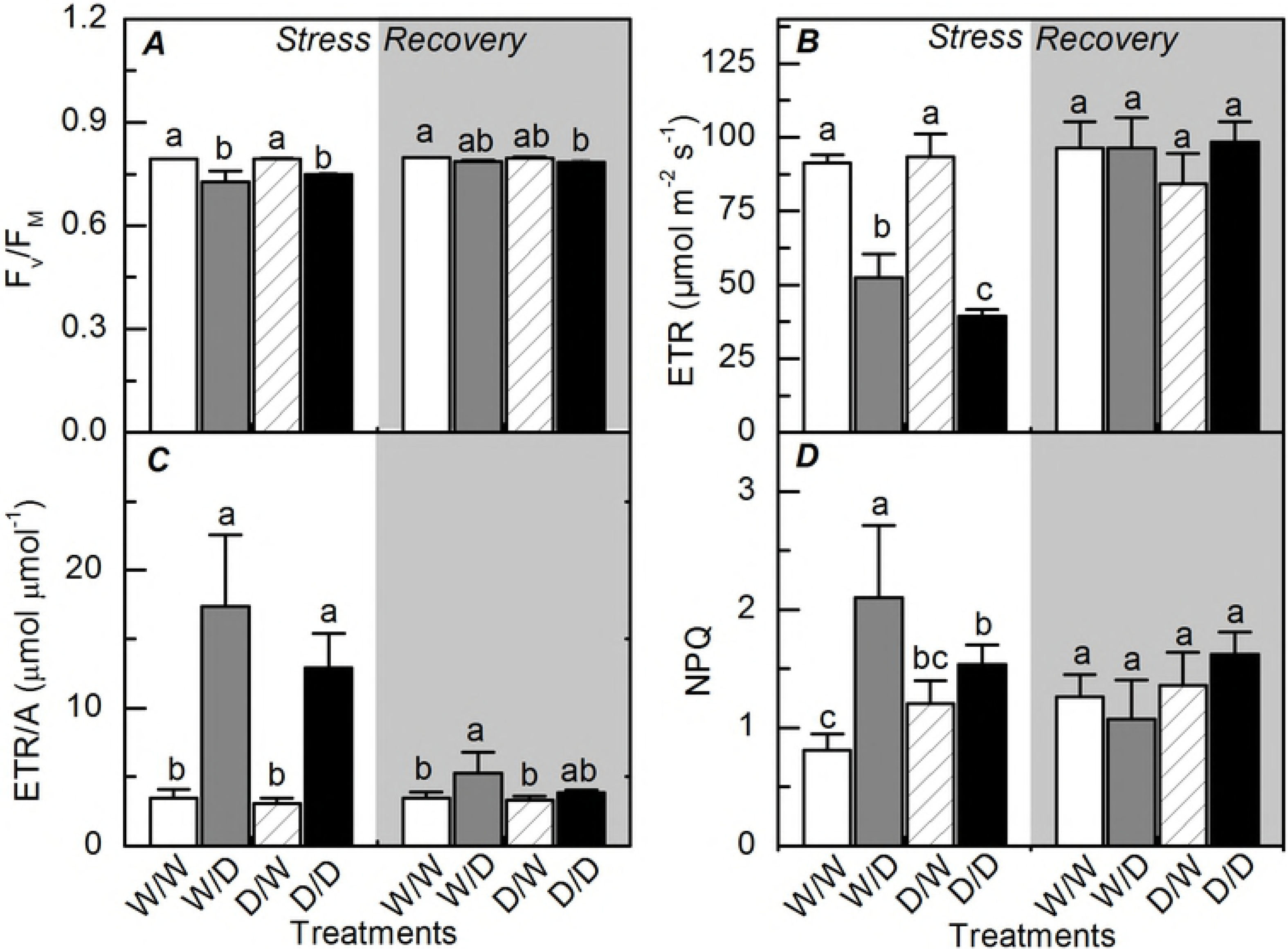
Photochemistry of daughter plants during and after water deficit. Potential quantum efficiency of photosystem II (*A*), the apparent electron transport rate estimated (*B*), ETR/A ratio (*C*), and the non-photochemical quenching of fluorescence (*D*) in sugarcane plants grown under well-watered conditions (W/W and D/W) or subjected to water deficit (W/D and D/D). Daughter plants were obtained from mother plants previously exposed to water deficit (D/W and D/D) or grown under well-watered conditions (W/W and W/D). Measurements were done during the water deficit (stress) and recovery (gray area) periods, as shown in Fig 1. Each histogram is the mean values + s.d. (n=4). Different letters mean statistical differences among treatments (p<0.05).

Leaf proline content was increased under water deficit and D/D plants presented the highest values. After the recovery period, W/D plants presented higher proline content than D/D plants (Fig 5).

**Fig 5.**
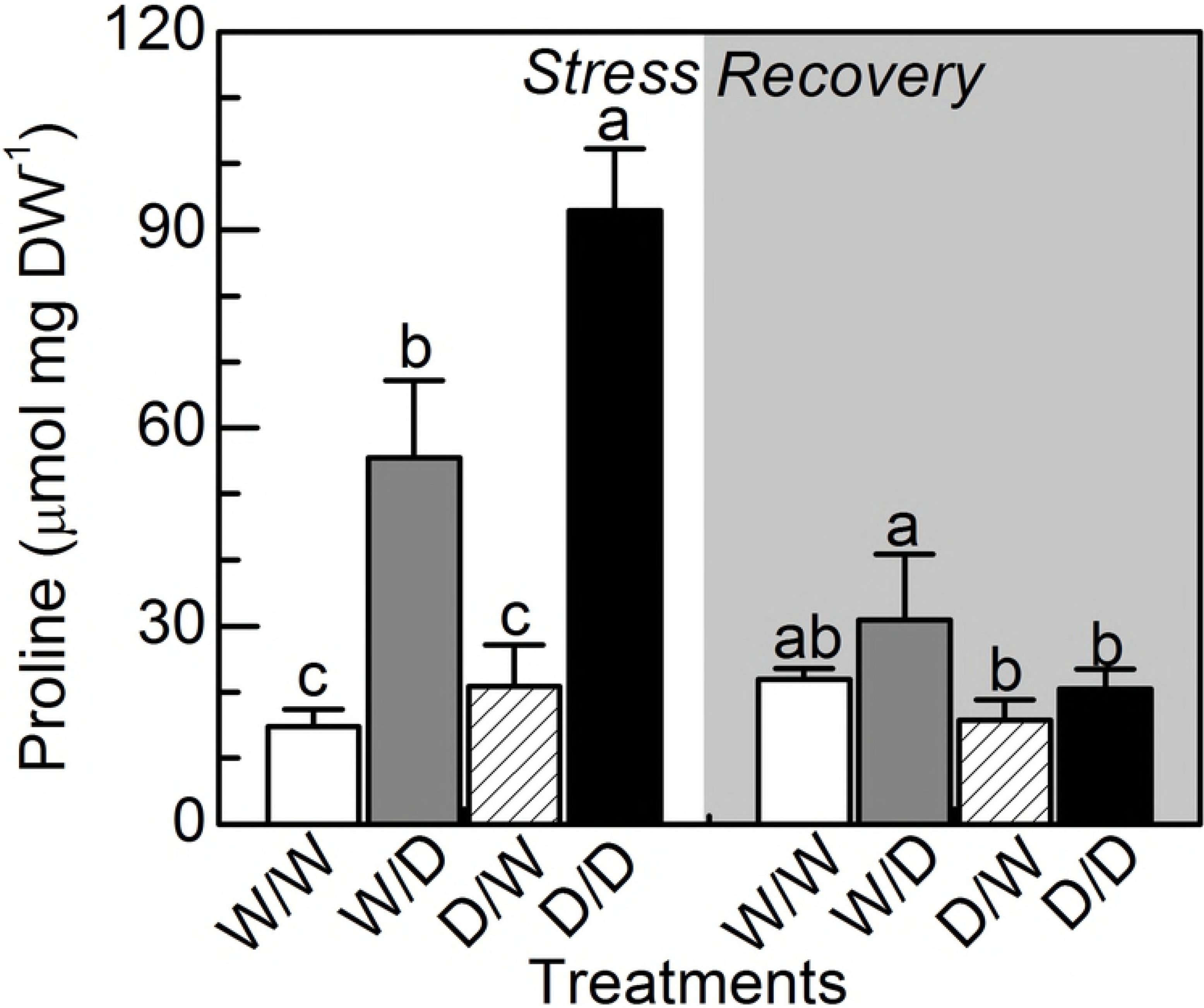
Leaf proline concentration in daughter plants under water deficit. Leaf proline concentration in sugarcane plants grown under well-watered conditions (W/W and D/W) or subjected to water deficit (W/D and D/D). Daughter plants were obtained from mother plants previously exposed to water deficit (D/W and D/D) or grown under well-watered conditions (W/W and W/D). Measurements were done during the water deficit (stress) and recovery (gray area) periods, as shown in Fig 1. Each histogram is the mean values + s.d. (n=4). Different letters mean statistical differences among treatments (p<0.05).

Leaf sucrose content was also increased by water deficit but only in plants originated from mother plants maintained under well-watered conditions, i.e. W/W vs. W/D (Fig 6A). Curiously, D/W plants had higher leaf sucrose content than W/W ones, suggesting an influence of mother plants. Such influence was also found in roots, with D/W plants presenting lower sucrose, soluble total sugars and total non-structural carbohydrates than W/W plants (Fig 6). Reductions in root concentrations of sucrose, soluble total sugars and total non-structural carbohydrates due to water deficit were found only in plants obtained from those ones that did not face drought (Fig 6E-H). When considering the total amount of non-structural carbohydrates in plants (Figure 6I), D/W plants had higher values than W/W plants and the carbohydrate partitioning between leaves (86% to 91%) and roots (9% to 15%) was similar among treatments (Fig 6J).

**Fig 6.**
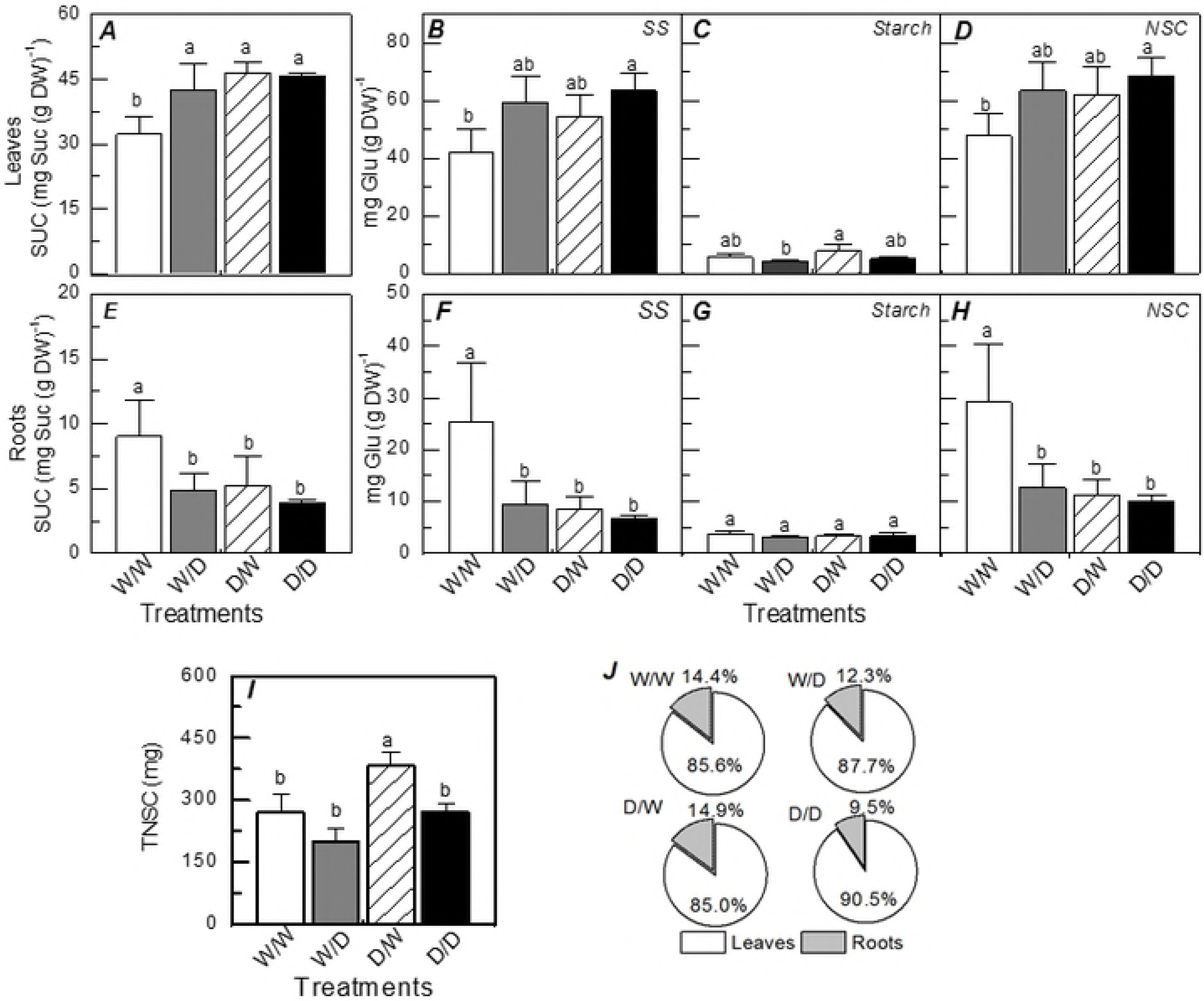
Leaf and root carbohydrates and their partitioning in daughter plants under water deficit. Sucrose (*A, E*) soluble sugars (*B, F*), starch (*C, G*) and non-structural carbohydrates (D, *H)*in leaves (A-D) and roots (E-H), amount of total non-structural carbohydrates in the entire plant (/) and their partitioning among plant organs (*J*) in sugarcane plants grown under well-watered conditions (W/W and D/W) or subjected to water deficit (W/D and D/D). Daughter plants were obtained from mother plants previously exposed to water deficit (D/W and D/D) or grown under well-watered conditions (W/W and W/D). Measurements were taken after 9 days of water deficit (maximum water deficit). Each histogram is the mean values + s.d. (n=4). Different letters mean statistical differences among treatments (p<0.05).

Regarding the antioxidant metabolism, leaf SOD and CAT activities were not affected either by water regime or plant origin (Fig 7A,D), while leaf H_2_O_2_ concentration and leaf APX activity increased due to water deficit (Fig 7B,C). The highest leaf APX activity was found in W/D plants (Fig 7C). In roots, non-significant changes were found for SOD and APX activities (Fig 7E,G). Root H_2_O_2_ concentration and CAT activity increased due to water deficit in daughter plants originated from well-watered mother plants (Fig 7F,H). On the other hand, root H_2_O_2_ concentration was reduced and root CAT activity did not change under water deficit when considering daughter plants originated from plants grown under cycles of water deficit (Fig 7F,H). Interestingly, D/W plants had higher root H_2_O_2_ concentration and higher root CAT activity than W/W plants (Fig 7F,H).

**Fig 7.**
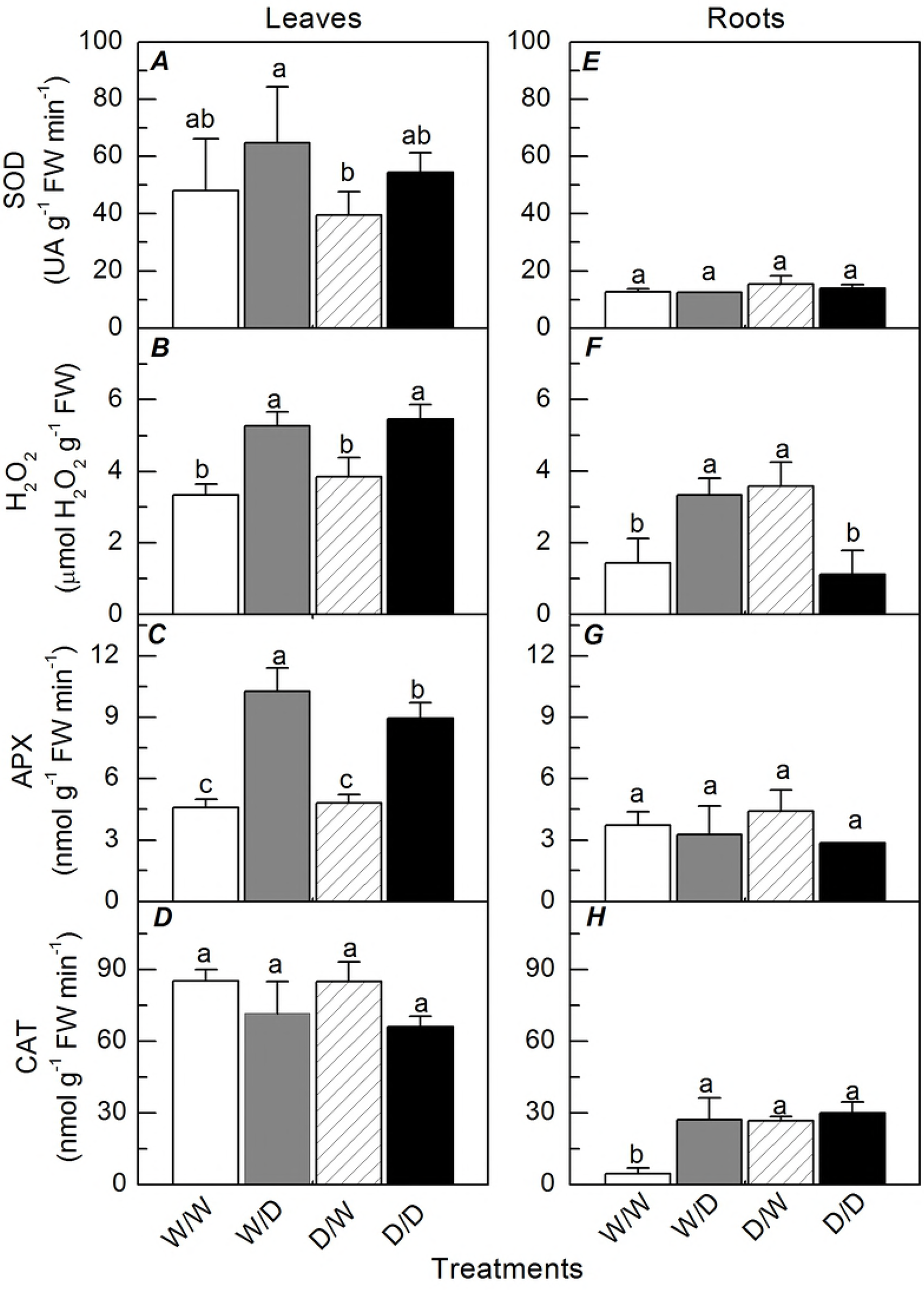
Antioxidant metabolism in daughter plants under water deficit. Activities of SOD (*A, E*), APX (*C, G*), CAT (*D, H*) and H_2_O_2_ concentration (*B, F*) in leaves (*A-D*) and roots (*E-H*) of sugarcane plants grown under well-watered conditions (W/W and D/W) or subjected to water deficit (W/D and D/D). Daughter plants were obtained from mother plants previously exposed to water deficit (D/W and D/D) or grown under well-watered conditions (W/W and W/D). Measurements were taken after 9 days of water deficit (maximum water deficit). Each histogram is the mean values + s.d. (n=4). Different letters mean statistical differences among treatments (p<0.05).

Water deficit reduced shoot biomass production regardless plant origin, but D/D plants had higher shoot biomass than W/D plants (Fig 8A). While daughter plants obtained from well-watered mother plants presented increases in root biomass under water deficit, the opposite was found in daughter plants obtained from mothers that experienced cycles of water deficit (Fig 8B). In general, root biomass of D/W plants was about four times higher than one of W/W plants, with D/D plants showing similar root biomass as compared to W/D plants. Leaf area was also reduced by water deficit (Fig 8C), but D/D plants had higher leaf area than daughter plants obtained from well-watered mothers, despite the water regime.

**Fig 8.**
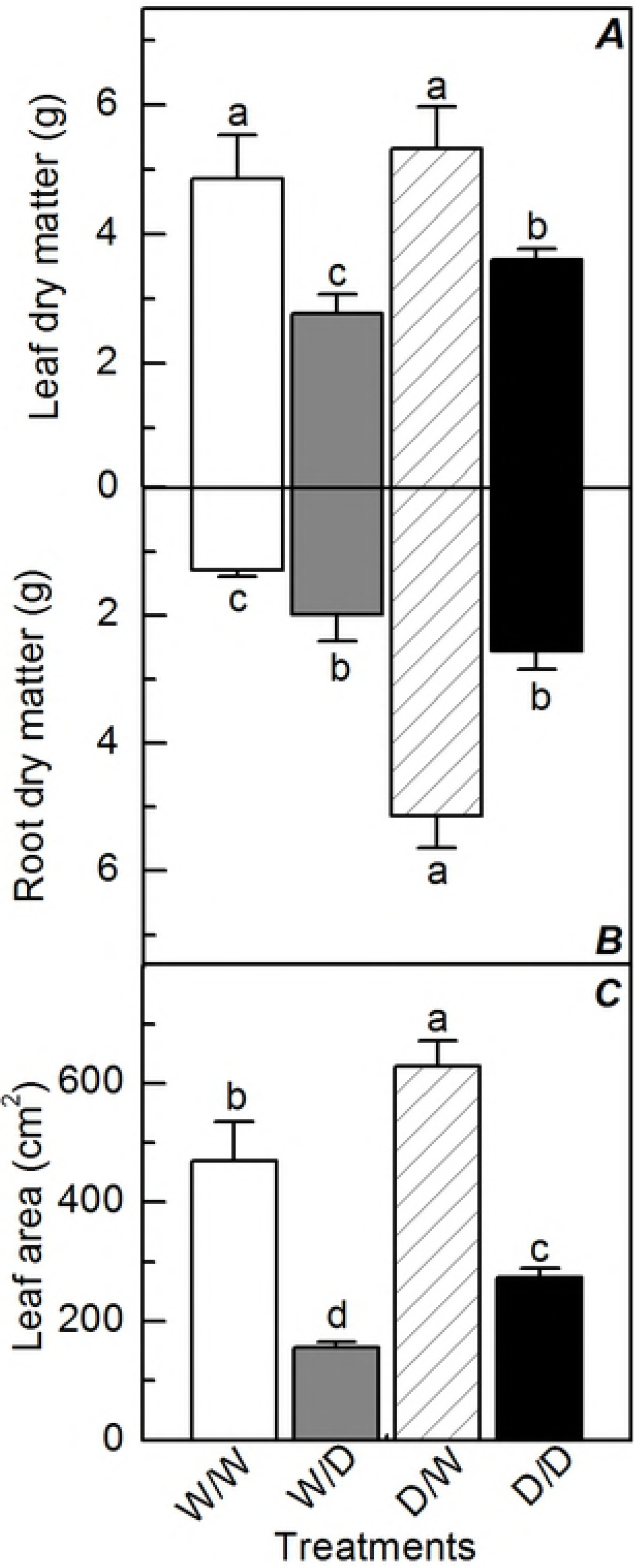
Biomass accumulation by daughter plants under water deficit. Leaf (A) and root (B) dry matter and leaf area (C) in sugarcane plants grown under well-watered conditions (W/W and D/W) or subjected to water deficit (W/D and D/D). Daughter plants were obtained from mother plants previously exposed to water deficit (D/W and D/D) or grown under well-watered conditions (W/W and W/D). Measurements were taken at the end of experiment. Each histogram is the mean values ± s.d. (n=4). Different letters mean statistical differences among treatments (p<0.05).

## Discussion

Herein, we induced transgenerational stress memory through vegetative propagation of sugarcane by inducing cycles of water deficit to mother plants. Epigenetic changes caused by varying environmental conditions help plants to adapt and have advantageous growth and acclimation under unstable environments [13]. Such epigenetic changes may manifest in the future generations, a transgenerational stress memory if mother plants were previously stressed [14]. Although sugarcane propagation does not involve meiotic recombination, mitotic alterations during vegetative propagation may also produce a source of epigenetic variation that helps plants to persist and succeed in environmental colonization [13].

Our findings indicate that propagules obtained from plants growing in areas with low water availability would be more tolerant to drought as compared to propagules of the same genotype grown under irrigation of in areas without occurrence of water deficit. Interestingly, daughters of mother plants that faced water deficit produced more biomass than ones from mother plants maintained well-watered, regardless water regime (Fig 8). This suggest that plants have increased their efficiency in using natural resources such as water and sunlight through the transgenerational stress memory. We have shown recently that stress memory is induced in sugarcane plants after three cycles of water deficit, with plants showing higher photosynthesis and improved growth under water limiting conditions [5]. Those previous results together with ones reported herein indicate that drought tolerance of sugarcane could be improved by water management and by selecting the propagation material for planting new crop fields.

When exposing mother plants to water deficit, stress memory was induced and the information likely stored in bud meristems, as suggested by improved performance of daughter plants obtained by vegetative propagation. Besides causing decreases in photosynthesis (S1 Fig) and biomass production (S2 Fig; S1 Table), cycles of dehydration and rehydration are able to create a number of chemical signals, such as increases in concentration of abscisic acid (ABA), a hormone that alter the expression pattern of many genes linked to drought response [31]. Changes in gene expression patterns might be stored through epigenetic changes such as DNA methylation and acetylation and induce stress memory [32]. In spite of a large decrease in biomass production of mother plants under water deficit (S2 Fig; S1 Table), daughter plants had faster sprouting and higher biomass than ones obtained from well-hydrated mother plants (Fig 8). Such improved plant growth due to the transgenerational stress memory was reported previously and it is likely linked to changes in DNA methylation [8], a research topic that should be further investigated for revealing the molecular bases of stress memory and tolerance in sugarcane.

The ability of clone plants in recovering the stored environmental information [15] can explain both morphological and physiological responses of D/D plants. D/D plants exhibited higher photosynthesis than W/D plants at recovery and this was caused by higher instantaneous carboxylation efficiency (Fig 1A,C). Regarding primary photochemistry, non-photochemical quenching was lower in D/D plants than in W/D plants, indicating less dissipation of energy as heat in the former ones (Fig 4D). Another interesting index suggesting stress memory is the water use efficiency [31], which indicates an optimization of CO_2_ assimilation per unit of H_2_O transpired in D/D plants under water limiting conditions (Fig 2C).

Interestingly, D/D plants were able to maintain metabolic activity and produce more biomass than W/D plants (Fig 8) even presenting lower leaf water potential (Fig 3A). As RWC was similar in W/D and D/D plants (Fig 3), our data indicate the occurrence of more intense osmotic adjustment in D/D plants. This can be explained by higher concentration of proline in leaves (Fig 5), an osmotic and osmoprotectant molecule [33]. During stressful conditions, high proline levels in D/D plants suggest that these plants have synthesized this osmolyte for adjusting the osmotic equilibrium and cell homeostasis, one form of memory according to [34]. After rehydration, there was a large degradation of proline in D/D plants, which would increase the remobilization of nitrogen to assimilatory pathways for resuming plant growth. Evidence of transgenerational stress memory was found even at the last day of rehydration, when D/D plants had higher photosynthesis (25.7±2.7 vs. 15.7±3.8 p,mol m^−2^ s^−1^) and integrated water use efficiency (7.2±0.3 vs. 6.3±0.4 µmol mol^−1^) than W/D plants.

Plants respond to abiotic stresses by altering their metabolism and accumulating substances such as sugars, amino acids and other metabolites with important roles in stress tolerance [35]. Maintenance of high sucrose concentration even under well-watered conditions may be another evidence of stress memory [36], as found in D/W plants (Fig 6A). In addition, plants obtained from mother plants that faced drought did not present any change in both leaf and root sucrose concentrations under water deficit (Fig 6A,E). Sucrose accumulation would help plants under water deficit by improving osmoregulation and protecting proteins and then maintaining photosynthesis under low water availability.

Low concentrations of ROS in plants previously exposed to stressful conditions could be an indication of stress memory [37]. However, our data indicate that exposure of mother plants to water deficit caused higher root H_2_O_2_ concentration in plants maintained under well-watered conditions (Fig 7F). In addition to its role in plant signaling [38], ROS accumulation is also associated with modifications in DNA methylation pattern [39], an epigenetic change that would store information and induce faster stress response. The presence of ROS in controlled amounts is important for plant growth, with plants showing higher H_2_O_2_ concentration in the region of root elongation [37]. In this way, high root H_2_O_2_ concentration in D/W plants (Fig 7F) would explain high root biomass (Fig 7F and 8B). In fact, H_2_O_2_ is produced by mitochondria during the synthesis of NADH and ATP for supplying aerobic plant metabolism in active growing regions [40].

Based on results reported herein, the next step towards the improvement of drought tolerance in sugarcane plants would be the evaluation of field-grown plants, considering the persistence of stress memory and its consequences for crop yield and biomass production as well as the genotypic variation within *Saccharum* complex.

## Conclusion

Our findings clearly show that sugarcane growth is improved in daughter plants obtained from mother plants that faced water deficit. The bases of such transgenerational stress memory should be further studied taking into account possible epigenetic markers. Our data also revealed that bud meristems of sugarcane are able to store information acquired from previous stressful events. Accumulation of H_2_O_2_ in roots is a possible chemical signal related to stress memory, being associated with improved root growth in well-watered plants. Benefits of such stress memory were noticed in leaf gas exchange and plants showed improved photosynthetic water use efficiency and faster recovery of photosynthesis after rehydration. As consequence, daughter plants obtained from stressed mothers exhibited improvements in biomass production, regardless of water conditions. Finally, our results bring a new perspective for the management of sugarcane fields as plant performance could be improved under field conditions due to a large root system and faster recovery of photosynthesis after facing water shortage.

## Author contributions

Conception and experimental design - Fernanda C C Marcos and Rafael V Ribeiro; Data collection - Fernanda C C Marcos, Neidiquele M Silveira and Paulo E R Marchiori; Data analysis and interpretation - Fernanda C C Marcos, Eduardo C Machado, Gustavo M Souza and Rafael V Ribeiro; Drafting of the article - Fernanda C C Marcos and Rafael V Ribeiro; Critical revision and final approval of the article - all authors.

## Conflict of interest statement

The authors declare that this research was conducted in the absence of any commercial or financial relationships that could be construed as a potential conflict of interest.

## Supporting information

**S1 Fig. Time course of leaf gas exchange in mother plants under water deficit.** Leaf CO_2_ assimilation of mother-plants maintained well-watered (W) or subjected to three cycles of water deficit (D). The grey area represents water withholding (nine days) and the dotted line indicates null photosynthesis. Each symbol represents the mean values ± s.d. (n = 4).

**S2 Fig. General view of mother plants after water deficit.** Visual aspect of mother plants grown under cycles of water deficit (left) or under well-watered conditions (right).

**S1 Table. Biomass accumulation by daughter plants under water deficit.** Biometry of mother plants grown under well-watered (reference) conditions or subjected to cycles of water deficit. Measurements were taken after 80 days of treatment. Different letters mean statistical differences between treatments (p<0.05).

## References

1. Ribeiro, RV, Machado, RS, Machado, EC, Machado, DFSP, Magalhães filho, JR, Landell, MGA. Revealing drought-resistance and productive patterns in sugarcane genotypes by evaluating both physiological responses and stalk yield. Experimental Agriculture. 2013; 49, 212–224. https://doi.org/10.1017/S0014479712001263

2. Sales, CRG, Marchiori, PER, Machado, RS, Fontenele, AV, Machado, EC, Silveira, JAG, Ribeiro, RV. Photosynthetic and antioxidant responses to drought during the sugarcane ripening? Photosynthetica. 2015; 53, 547–554. https://doi.org/10.1007/s11099–015–0146-x

3. Bruce, TJA, Matthes, MC, Napier, JA, Pickett, JA. Stressful “memories” of plants: Evidence and possible mechanisms. Plant Science. 2007; 173, 603–608. https://doi.org/10.1016/j.plantsci.2007.09.002

4. Galle, A, Florez-Sarasa, I, El Aououad, H, Flexas, J. (2011). The Mediterranean evergreen *Quercus ilex and the semideciduous Cistus albidus* differ in their leaf gas exchange regulation and acclimation to repeated drought and re-watering cycles. Journal of Experimental Botany. 2011; 62, 5207–5216. https://doi.org/10.1093/jxb/err233

5. Marcos, FCC, Silveira, NM, Mokochinski, JB, Sawaya, ACHF, Marchiori, PER, Machado, EC, Souza, GM, Landell, MGA, Ribeiro, RV. Drought tolerance of sugarcane is improved by previous exposure to water deficit. Journal of Plant Physiology. 2018; 223, 9–18. https://doi.org/10.1016/j.jplph.2018.02.001

6. Walter, J, Jentsch, A, Beierkuhnlein, C, and Kreyling, J. Ecological stress memory and cross tolerance in plants in the face of climate extremes. Environmental and Experimental Botany. 2013; 94, 3–8. https://doi.org/10.1016/j.envexpbot.2012.02.009

7. Izanloo, A, Condon, AG, Langridge, P, Tester, M, Schnurbusch, T. Different mechanisms of adaptation to cyclic water stress in two South Australian bread wheat cultivars. Journal of Experimental Botany. 2008; 59, 3327–3346. https://doi.org/10.1093/jxb/ern199

8. Hauser, MT, Aufsatz, W, Jonak, C, Luschnig, C. Transgenerational epigenetic inheritance in plants. Biochim Biophys Acta. 2011; 1809, 459–468. https://doi.org/10.1016/j.bbagrm.2011.03.007

9. Ding, Y, Fromm, M, Avramova, Z. Multiple exposures to drought “train” transcriptional responses in *Arabidopsis*. Nature Communication. 2012; 3, 740. https://doi.org/10.1038/ncomms1732

10. Thellier, M, Lüttge, U. Plant memory: a tentative model. Plant Biology. 2012; 15, 112. https://doi.org/10.1111/j.1438-8677.2012.00674.x

11. Chinnusamy, V, Zhu, JK. Epigenetic regulation of stress responses in plants. Current Opinion in Plant Biology. 2009; 12, 133–139. https://doi.org/10.1016/j.pbi.2008.12.006

12. Boyko, A, Kovalchuk, I. Genome instability and epigenetic modification-heritable responses to environmental stress? Current Opinion in Plant Biology. 2011; 14, 260–266. https://doi.org/10.1016/j.pbi.2011.03.003

13. Dodd, R, Douhovnikoff, V. Adjusting to Global Change through Clonal Growth and Epigenetic Variation. Frontiers in Ecology and Evolution. 2016; 4, 86. https://doi.org/10.3389/fevo.2016.00086

14. Boyko, A, Blevins, T, Yao, Y, Golubov, A, Bilichak, A, Ilnytskyy, Y, Hollander, J, Meins, Jr F, Kovalchuk, I. Transgenerational adaptation of Arabidops is to stress requires DNA methylation and the function of dicer-like proteins. PlosONE. 2010; 5, e9514. https://doi.org/10.1371/journal.pone.0009514

15. Latzel, V, Rendina González, AP, Rosenthal, J. Epigenetic Memory as a Basis for Intelligent Behavior in Clonal Plants. Front Plant Science. 2016; 7, 1354. https://doi.org/10.3389/fpls.2016.01354

16. Dos Santos, HG. Sistema Brasileiro de Classifiçãgao de solos. 3th ed. Brasília, DF.: Embrapa; 2013.

17. Dias, FL, Rosseto, R. Calageme adubação da cana-de-açúcar. In: Atualização em produção de cana-de-açúcar, ed Segato, SV, Pinto, AS Jendiroba, Nóbrega, JCM. Piracicaba: Livro Ceres, 2016. pp. 107–119.

18. Sarruge, JR. Soluções nutritivas. Summa Phytopathologica. 1975; 1, 231–233. http://www.scielo.br/scielo.php?script=sci_nlinks&ref=000052&pid=S0071-1276197800010001700009&lng=es

19. Edwards, GE, Baker, NR. Can CO_2_ assimilation in maize leaves be predicted accurately from chlorophyll fluorescence analysis? Photosynth Research. 1993; 37, 89–102. https://doi.org/10.1007/BF02187468

20. Baker NR. Chlorophyll Fluorescence: A probe of photosynthesis: *in vivo*. Annual Review of Plant Biology. 2008; 59, 89–113. https://doi.org/10.1146/annurev.arplant.59.032607.092759

21. Weatherley, PE. Studies in the water relations of the cotton plant. I. The field measurement of water deficits in leaves. New Phytologist. 1950; 49, 81–87. https://doi.org/10.1111/j.1469–8137.1950.tb05146.x

22. Bieleski, RL, Turner, A. Separation and estimation of amino acids in crude plant extracts by thinx’-layer electrophoresis and chromatography. Analytical Biochemistry. 1966; 17, 278–293. https://doi.org/10.1016/0003-2697(66)90206–5

23. Dubois, M, Gilles, KA, Hamilton, JK, Rebers, PA, Smith, F. Colorimetric method for determination of sugars and related substances. Analytical Biochemistry. 1956; 28, 350–356. https://doi.org/10.1021/ac60111a017

24. Van Handel, E. Direct microdetermination of sucrose. Analytical Biochemistry. 1968; 22, 280–283. https://doi.org/10.1016/0003–2697(68)90317–5

25. Amaral, LIV, Gaspar, M, Costa, PMF, Aidar, MPM, Buckeridge, MS. Novo metodo enzimatico rapido e sensfvel de extragao e dosagem de amido em materiais vegetais. Hoehnea. 2007; 34, 425–431. http://dx.doi.org/10.1590/S2236–89062007000400001

26. Rena, AB, Masciotti, GZ. The effect of dehydration on nitrogen metabolism and growth of bean cultivars *(Phaseolus vulgaris* L.). Ceres. 1976; 23, 288–301. http://www.scielo.br/scielo.php?script=sci_nlinks&ref=000041&pid=S0006-8705198100010000500010&lng=en

27. Alexieva V, Sergiev I, Mapelli S, and Karanov E. The effect of drought and ultraviolet radiation on growth and stress markers in pea and wheat. Plant Cell Environment 2001; 24, 1337–1344. https://doi.orq/10.1046/j.1365-3040.2001.00778.x

28. Giannopolitis, O, Ries SK. Superoxide dismutase: I. Occurrence in higher plants. Plant Physiology. 1977; 59, 309–14. https://doi.org/10.1104/pp.59.2.309

29. Havir, EA, McHale NA. Biochemical and development characterization of multiples forms of catalase in Tobacco-leaves. Plant Physiology. 1987; 84, 450–455. https://doi.org/10.1104/pp.84.2.450

30. Nakano, Y, Asada, K. Hydrogen peroxide is scavenged by ascorbate-specific peroxidase in spinach chloroplasts. Plant Cell Physiology. 1981; 22, 867–880. https://doi.org/10.1093/oxfordiournals.pcp.a076232

31. Fleta-Soriano, E, Pintó-Marijuan, M, Munné-Bosch, S. Evidence of Drought Stress Memory in the Facultative CAM, *Aptenia cordifolia:* Possible Role of Phytohormones. PlosONE. 2015; 10, e0135391. https://doi.org/10.1371/iournal.pone.0135391

32. Avramova, Z. Transcriptional ‘Memory’ of a Stress; transient chromatin and memory (epigenetic) marks at stress response genes. The Plant Journal. 2015; 83, 149–159. https://doi.org/10.1111/tpi.12832

33. Szabados, L, Savouré A. Proline: a multifunctional amino acid. Trends Plant Science. 2010; 15, 89–97. https://doi.org/10.1016/j.tplants.2009.11.009

34. Ding, Y, Liu, N, Virlouvet, L, Riethoven, JJ, Fromm, M, Avramova, Z. Four distinct types of dehydration stress memory genes in *Arabidopsis thaliana.* BMC Plant Biology. 2013; 13: 229. https://doi.org/10.1186/1471–2229–13–229

35. Verlues, PE, Agarwal, M, Katiyar-Agarwal, S, Zhu, J, Zhu, JK. Methods and concepts in quantifying resistance to drought, salt and freezing, abiotic stress that affect plant water status. Plant Journal. 2006; 45, 523–539. https://doi.org/10.1111/M365-313X.2005.02593.x https://doi.org/10.1111/M365-313X.2005.02593.x

36. Crisp, PA, Ganguly, D, Eichten, SR, Borevitz, J.O., Pogson, B.J. Reconsidering plant memory: Intersections between stress recovery, RNA turnover, and epigenetics. Science Advances. 2016; 2: e1501340. https://doi.org/10.1126/sciadv.1501340

37. Hu, T, Jinb, Y, Lia, H, Amomboa, E, Fua, J. Stress memory induced transcriptional and metabolic changes of perennial ryegrass (Lolium perenne) in response to salt stress. Physiologia Plantarum. 2015; 156, 54–69. https://doi.org/10.1111/ppl.12342

38. Foyer, CH, Noctor, G. Oxidant and antioxidant signalling in plants: a re-evaluation of the concept of oxidative stress in a physiological context. Plant Cell Environment. 2005; 28, 1056–1071. https://doi.org/10.1111/j.1365–3040.2005.01327.x

39. Peng, H, Zang, J. Plant genomic DNA methylation in response to stresses: Potential applications and challenges in plant breeding. Progress in Natural Science. 2009; 19, 1037–1045. https://doi.org/10.1016/j.pnsc.2008.10.014

40. Gill, SS, Tuteja, N. Reactive oxygen species and antioxidant machinery in abiotic stress tolerance in crop plants. Plant Physiology and Biochemistry. 2010; 48, 909–930. https://doi.org/10.1016/j.plaphy.2010.08.016

